# Feeding intensity and molecular prey identification of the common long-armed octopus, *Octopus minor* (Mollusca: Octopodidae) in the wild

**DOI:** 10.1101/707349

**Authors:** Qi-Kang Bo, Xiao-Dong Zheng, Zhi-Wei Chen

## Abstract

The common long-armed octopus, *Octopus minor*, is an important component of systems and supports the local fisheries in the coastal areas of northern China. And because of the overfishing, the national germplasm reserve was established as conservation area at the Moon Lake for the genetic resource of *O.minor* in 2012. For the fishery management and artificial breeding, especially for the management of exclusive conservation reserves, its role in the ecosystem requires assessment. Therefore, the feeding intensity of *O. minor* was studied from April to July 2014 when females reaching maturation, and prey composition was identified from stomach contents using a DNA barcoding method. Of the 172 sampled octopuses, 66 had stomach contents that were nearly digested into pulp. Maximum feeding intensity occurred during the month of April and the feeding intensity of the females was greater than that of the males in April and May. A considerable overall reduction of feeding intensity in both sexes occurred from April to July. A total of 8 species were identified as the prey of *O. minor*. Based on homology searches and phylogenetic analysis, of 60 sequences, 30 matched with fish (50.00%, by number), 13 with crustaceans (21.66%), one with annelid (1.66%), one with nematode (1.66%) and 15 with itself (25.00%). These results confirm that *O. minor* has habitual nature of strong dietary preference, with Gobiidae families (62.79%, by number) being an important prey during the time when females reach sexual maturation. From April to July, the observed cannibalism showed an increasing trend.

## Introduction

The common long-armed octopus, *Octopus minor* (Sasaki 1920) is a benthic and neritic octopod that has become a commercially important species in the north of China and in South Korea [1, 2] and is identified to be furthermore ecologically important as a generalist predator. Because of overfishing, a national germplasm reserve was established at the Moon Lake (Fig 1) in 2012 to serve as conservation for the genetic resource of *O. minor*. Moon Lake is a shallow (depth <3 m) lagoon with a muddy, bivalve-covered bottom, and patches of sea grass growing in the subtidal zone, which provides a good habitat for *O. minor*. Additionally, a program of artificial propagation and release of *O. minor* has been launched in the north of China [2].

**Fig 1.**
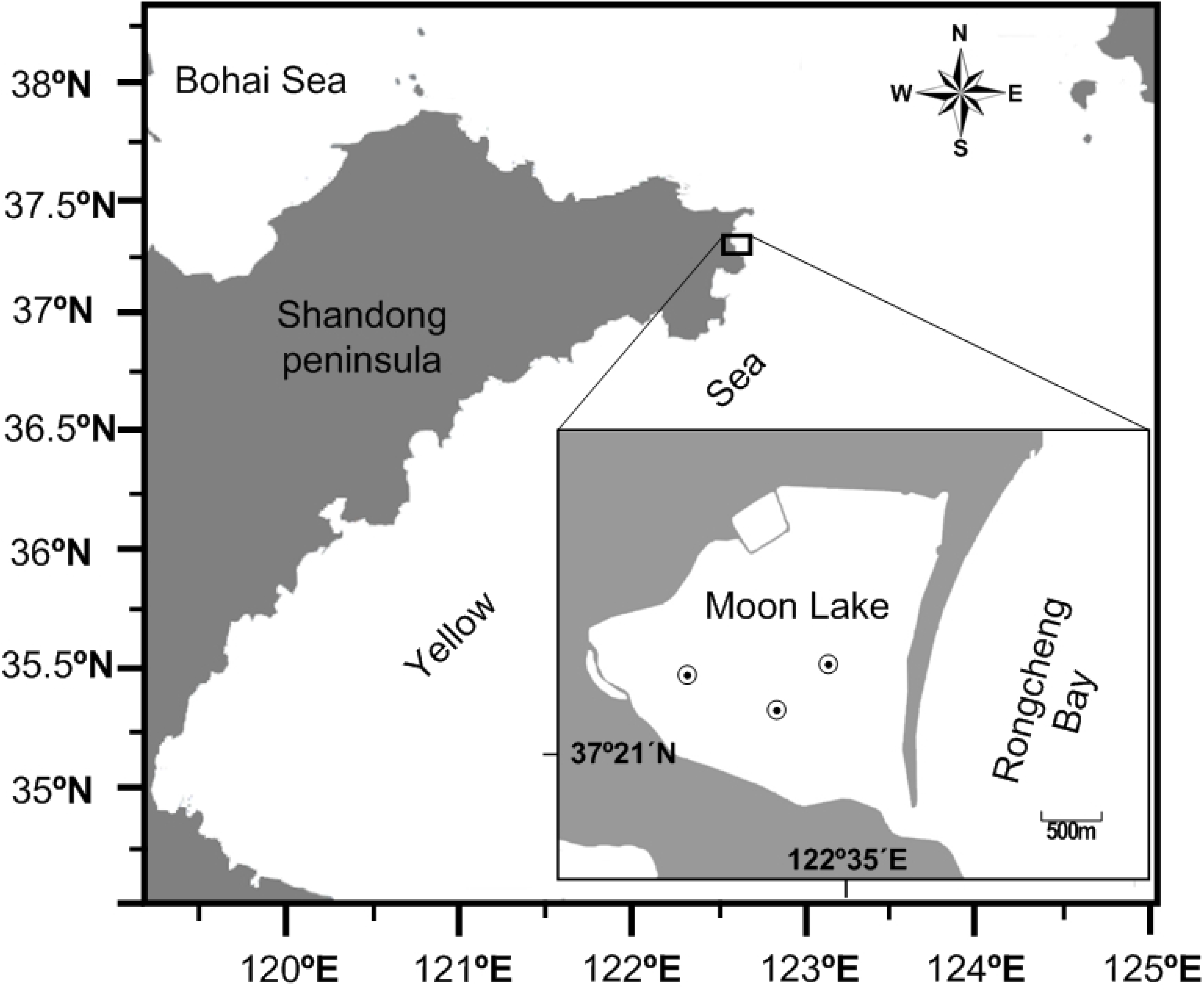
The location of the sampling site in Moon Lake and the dots show the sampling site.

The five methods used to assess the natural feeding habits of marine animals include the examination of prey remains from middens [3], direct observation [4], stomach content analysis [5, 6], isotopic assessment [7, 8], trophic tracing [9] and molecular prey identification [10–12]. In some studies, several different methods have been used in combination [11, 13, 14].

A range of factors has made it difficult to determine the natural diet of *O. minor*; the primary challenge that the oesophageal diameter of the octopus is physically limited as it passes through the brain; therefore, the octopus’ beak bites small pieces of tissue to swallow, avoiding the ingestion of hard skeletal material [11], and it is difficult to make an accurate identification of pray with lack of hard skeletal material. Moreover, it is difficult to be collected on a large scale. *O. minor* burrow deep (0.3-0.6 meter) interconnecting tunnels as nest in muddy marine bottom, leaving only digging-holes and breathing-holes on the surface, into which it hides itself [15]; their prey remains are left underground. Even if the prey remains were pushed out of the nests, the light material would be easily removed by biotic and abiotic factors, while the heavier material would be buried under the mud. Rapid digestion rates [16] and external ingestion [17] make the stomach contents visually unidentifiable. These specialized feeding strategies tend to bias data on prey species when morphological analysis is used.

Food is an important factor to the extent that it governs growth, fecundity and migratory movements. An understanding of the relationship between octopus species and their favorite food items helps to locate potential feeding grounds, which may, in turn, be helpful for the exploitation of these resources. The extent of the variation in feeding must be taken into account when studying the diet of a generalist predator. Octopods are opportunistic and their diets are often affected by the abundance of available prey [3, 18]. The species, size and type of their diet in the wild are often dominated by many factors such as maturity stage, seasonal variation [19], the body size which reflects the prey ability [3, 14] and benthic assemblages which determine prey availability [20]. In addition, what should be taken into account is that octopods exhibit strong dietary preferences when given the same opportunity to different diets items [18, 21–24].

Few studies have focused on the ecology of *O. minor*, whose diet is roughly known to include crustaceans, molluscs, polychaetes and fish [25, 26]. Investigation into the natural diet of this species during sexual maturation, are of more research value, and can provide trophic relationship information for fishery management, especially for the management of exclusive conservation reserves. It also provides a pray reference for aquaculture and artificial breeding of this commercially important specie. In this study, we applied the method of molecular prey identification, DNA barcoding method, to identify natural prey of *O. minor* based on stomach contents.

## Materials and methods

### Specimen collection

All the analyses have been carried out using freshly dead specimens collected from fishermen of the Mashan Group Co. Ltd. No use of live animals has been required for this study and no specific permissions were needed for the sampling activities in all of the investigated areas because our species of interest is commercially harvested (not endangered nor protected) and it was caught in areas where fishing is allowed. One hundred and seventy-two *O. minor* specimens were captured using traps placed during the evenings and collected at dawn from Moon Lake (122°35’E, 37°21’N, Fig 1) from April to July 2014. The same number of males and females were collected each month. All specimens were brought to the laboratory where their stomach contents were removed. Each stomach was opened and the contents were flushed into cryogenic vials. The stomach contents, potential residue of prey, were weighed and then preserved in 70% ethanol at −20°C for later DNA analysis [27]. Stomach content samples were numbered monthly, AP01 to AP19, MA01 to MA13, JN01 to JN19 and JL01 to JL15 from April to July, respectively. To avoid potential contaminants (e.g., blood and tissue attached to the stomach from the predator), the exterior surface of each stomach was washed with sterile, distilled water before removing the stomach contents [28]. The dorsal mantle length (ML), the distance between the posterior midpoint of the mantle and the midpoint of the eyes, and wet body weight (W) were also measured. This basic information and data were showed in Table 1. Feeding intensity during the study months was determined based on the degree of fullness of the stomach. The condition of feeding activity was determined from observations of the degree of stomach distension as described by Pillay (1952) [29]. Octopuses with stomachs that were gorged, full, ¾full, ½full were considered to have been actively feeding (AF), while stomach ¼full, little and empty were considered to denote poor feeding activity (PAF). The percentage of octopus in the AF and PAF condition in both sexes for each month was calculated.

**Table 1.** The sample number, stomach content specimens number (Num.), mantle length (ML, ±SD) and total weight (W, ±SD) of *O. minor* specimens collected each month from Moon Lake

| Sample date | Sample number | Female |  | Male |  | Num. |
| --- | --- | --- | --- | --- | --- | --- |
|  |  | ML (cm) | W (g) | ML (cm) | W(g) |  |
| 20/4/2014 | 52 | 8.68±0.95 | 120.47±24.99 | 8.60±0.89 | 135.65±28.26 | 19 |
| 14/5/2014 | 40 | 8.95±1.86 | 117.29±37.42 | 8.87±0.71 | 128.68±25.71 | 13 |
| 20/6/2014 | 40 | 10.55±1.14 | 192.46±40.28 | 10.30±1.09 | 202.88±42.66 | 19 |
| 16/7/2014 | 40 | 9.99±1.23 | 172.95±37.96 | 9.57±0.89 | 200.35±36.80 | 15 |

### DNA extraction and sequence acquisition

Stomach contents were evenly ground in a homogenizer. Duplicate 20-40 mg samples of mixed tissue items were subsampled from each stomach content and genomic DNA was then isolated by standard phenol-chloroform purification procedures. A fragment of the mitochondrial cytochrome oxidase I (CO I) gene was amplified using the universal primers [30].

Each polymerase chain reaction (PCR) was carried out in 50-μL volumes containing 2U Taq DNA polymerase (Takara Co.), about 60 ng template DNA, 0.2 mM dNTPs, 0.25 μM of each primer, 2 mM MgCl_2_ and 1×PCR buffer. The PCR amplification was performed on a GeneAmp^®^ 9700 PCR System (Applied Biosystems). Cycling conditions consisted of an initial denaturation at 94°C for 3 min, followed by 35 cycles of: denaturation at 94°C for 1 min, annealing at 50°C for 1 min, extension at 72°C for 1 min, and a final step of 5 min at 72°C.

Amplification products were confirmed by 1.5% TBE agarose gel electrophoresis stained with ethidium bromide. The cleaned product was prepared for sequencing using the BigDye Terminator Cycle Sequencing Kit (ver.3.1, Applied Biosystems) and sequenced bidirectionally using an ABI PRISM 3730 (Applied Biosystems) automatic sequencer. PCR products producing multiple bands indicated that more than one prey species were present. Those PCR products were cloned using the TOPO TA Cloning Kit (Invitrogen). Eight colonies per sample were selected for colony PCR amplification and sequencing using the primers M13 (forward): GTAAAACGACGGCCAG, andM13 (reverse): CAGGAAACAGCTATGAC.

### DNA analyses and statistical analysis

Sequences of CO I were assembled and edited separately using DNASTAR software (DNASTAR, Inc.), and then aligned with CLUSTAL_X 1.81 using the default settings [31]. Sequences were considered to be part of the same ‘‘operational taxonomic unit’’ (OTU), if there was less than a 1% sequence divergence, allowing for the intraspecific variation and Taq polymerase errors [11, 13].

All of the obtained sequences were identified using the Identification System (IDS) in the Barcode of Life Database (BOLD, www.boldsystems.org) and the Basic Local Alignment Search Tool (BLAST) query algorithm in GenBank to establish whenever possible the identification of the ingested material. The maximum likelihood (ML) method was chosen to infer evolutionary history. Bootstrap probabilities with 1,000 replications were calculated to assess reliability on each node of the ML tree. Sequence divergence calculation and evolutionary analyses were conducted in MEGA 6 software based on the Kimura 2-parameter model (K2P) [32]. The ML tree contained all of the sequences obtained from the stomach contents, together with the closest matches to each sequence that were downloaded from BOLD databases and GenBank. The criteria to assign identification to at the species level required the sequence similarity display >98% in the BOLD database and BLAST [11] and, if not, identification was restricted to the highest taxonomic lineage supported by bootstrap probabilities higher than 70% in the consensus tree [14, 15]. The AF and PAF differences the between sexes were verified by the Chi-squared fit test (χ^2^). Statistical analyses were performed using IBM SPSS statistics version 20 (IBM, Chicago, IL, USA). To corroborate that the number of analyzed stomachs was adequate for diet description, a cumulative prey curve was generated using the Estimate S Version 8.2 based on the prey identified [33]. The number of samples was assumed to be sufficient to describe the diet when the curve approaches the asymptote.

## Results

### Feeding intensity

Of the 172 octopus individuals, 66 had contents in their stomachs (Table 1) and feeding intensity of *O. minor* varied to an extent in respect to the months, and more than half of the stomachs were found to be empty (Table 2). There was significant difference in the feeding activity between July and other months (p<0.05) and no difference among the first three months (p>0.05). However, the AF percentage of octopus from April is greater than that of other months, meaning the maximum feeding intensity occurred during the month of April. A considerable overall reduction in the intensity of feeding for both sexes occurred from April to July (Table 2). The feeding intensity of the females is greater than that of the males based on the percentage of AF and PAF between the sexes in each month, and in April and May the feeding intensity of the females was significantly greater than that of the males (Table 3). That is, compared to the males, more female individuals were actively feeding.

**Table 2.** The distribution of stomach with different degree of distension and the percentage of octopus with active feeding activity (AF) from April to July. Data in the same column having different superscripted letters are significantly different (p<0.05).

| Month | Stomach status (%) |  |  |  |  |  |  | Feeding activity (%) |
| --- | --- | --- | --- | --- | --- | --- | --- | --- |
| | Gorge<br>d | Full | $\frac{3}{4}$Full | $\frac{1}{2}$Full | $\frac{1}{4}$Full | Little | Empty | AF |
| Apr. | 5.77 | 5.77 | 5.77 | 13.46 | 1.92 | 3.85 | 63.46 | 30.81<sup>a</sup> |
| May. | 6.00 | 16.00 | 5.00 | 2.50 | 3.00 | 0.00 | 67.50 | 29.50<sup>a</sup> |
| Jun. | 5.00 | 7.50 | 7.50 | 10.00 | 17.50 | 0.00 | 52.50 | 30.00<sup>a</sup> |
| Jul. | 0.00 | 0.00 | 5.00 | 5.00 | 17.50 | 10.00 | 62.50 | 10.00<sup>b</sup> |

**Table 3.**
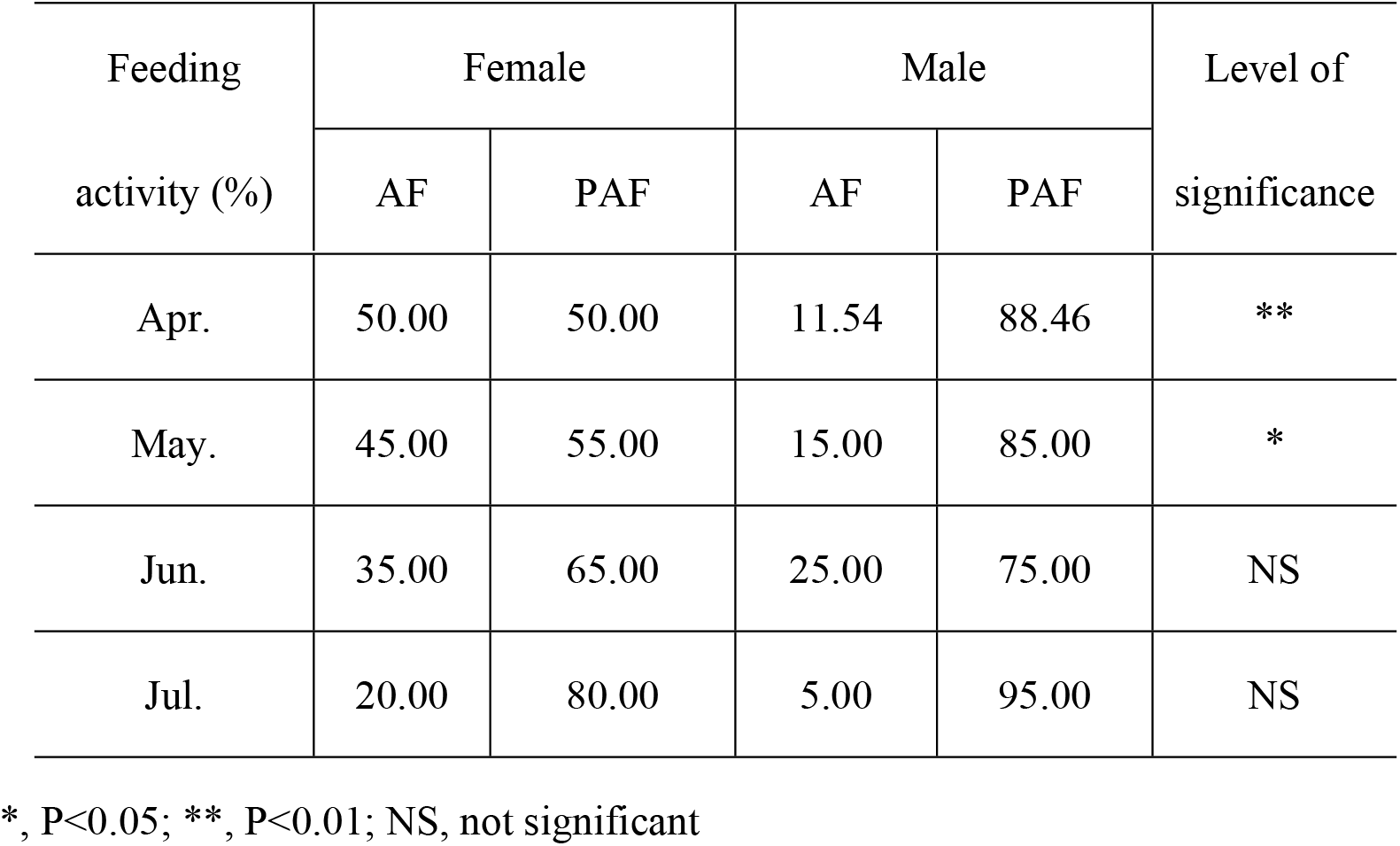
The percentage of octopus with active feeding activity (AF) and poor feeding activity (PAF) in both sexes from April to July, including the difference of feeding intensity between females and males in each month by Chi square fit test.

| Feeding<br>activity (%) | Female |  | Male |  | Level of<br>significance |
| --- | --- | --- | --- | --- | --- |
|  | AF | PAF | AF | PAF |  |
| Apr. | 50.00 | 50.00 | 11.54 | 88.46 | ** |
| May. | 45.00 | 55.00 | 15.00 | 85.00 | * |
| Jun. | 35.00 | 65.00 | 25.00 | 75.00 | NS |
| Jul. | 20.00 | 80.00 | 5.00 | 95.00 | NS |

\*, P<0.05; \*\*, P<0.01; NS, not significant

### Molecular prey identification

All stomach contents were nearly digested into pulp and as a result, the prey items were impossible to visually identify. A total of 59 stomach contents yielded amplifiable DNA and 60 sequences were obtained, ranging from 500bp to 708 bp (Table 4). All sequences were submitted to GenBank (Accession numbers MK688462-MK688521). Six OTUs were established, with a maximum sequence divergence of 0.1%. Of the 60 sequences, 59 clones showed similarities higher than 98% to reference sequences, allowing identification at species level. Only one clone displayed 97.56% similarity to reference sequences, and it was assigned to the *Diopatra* genus level (Fig 2). One stomach contained two kinds of prey, and the remaining 58 samples contained only one kind. The duplicate parallel PCR products from the contents of each stomach yielded the same sequences.

**Table 4.**
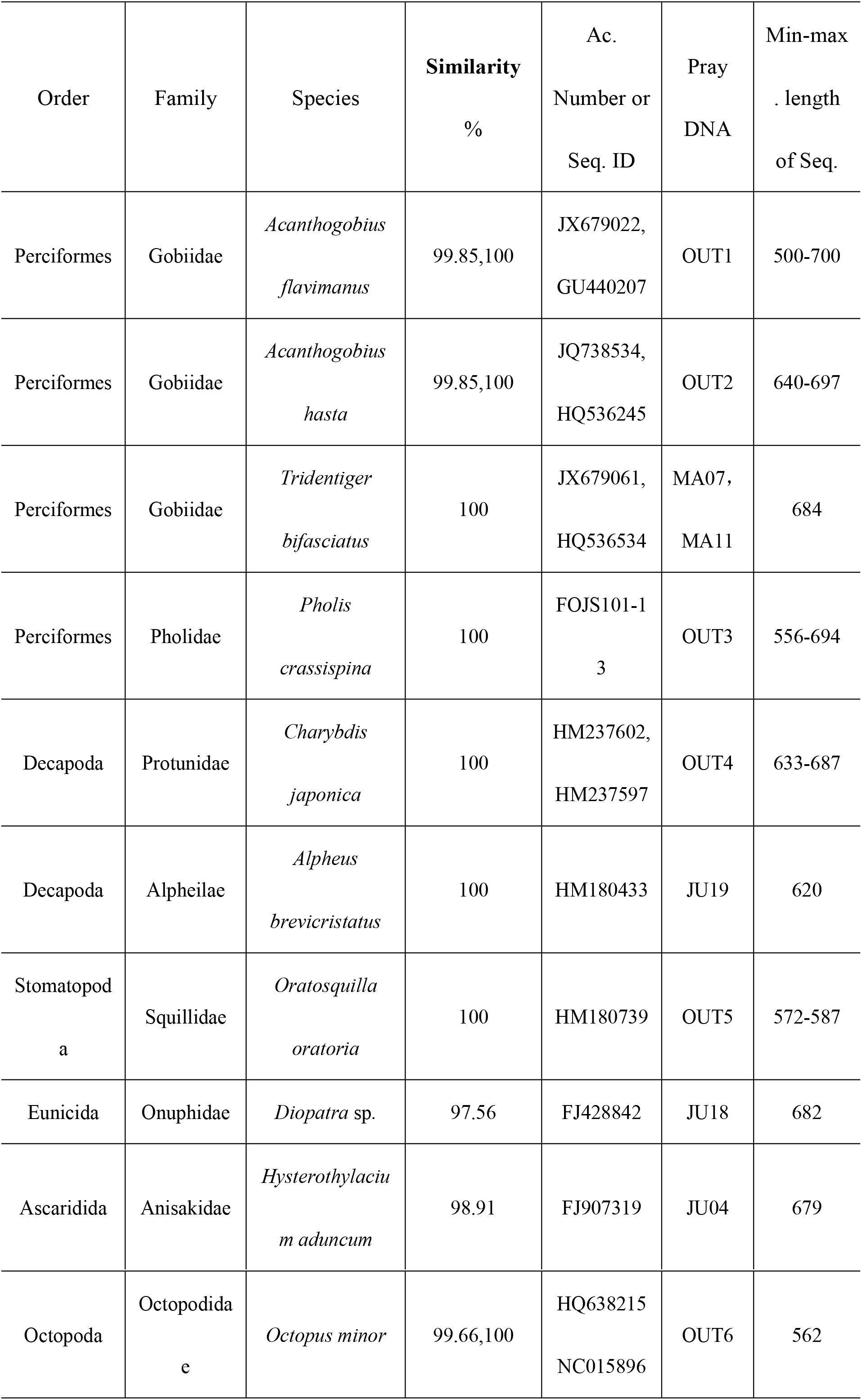
Pray DNA detected in wild *O. minor* by cloning the CO I fragment gene, including number per month and min-max length of sequences, GenBank Accession numbers and Sequence ID of closest matches, percentages of similarity obtained from BLAST and BOLD.

**Fig 2.**
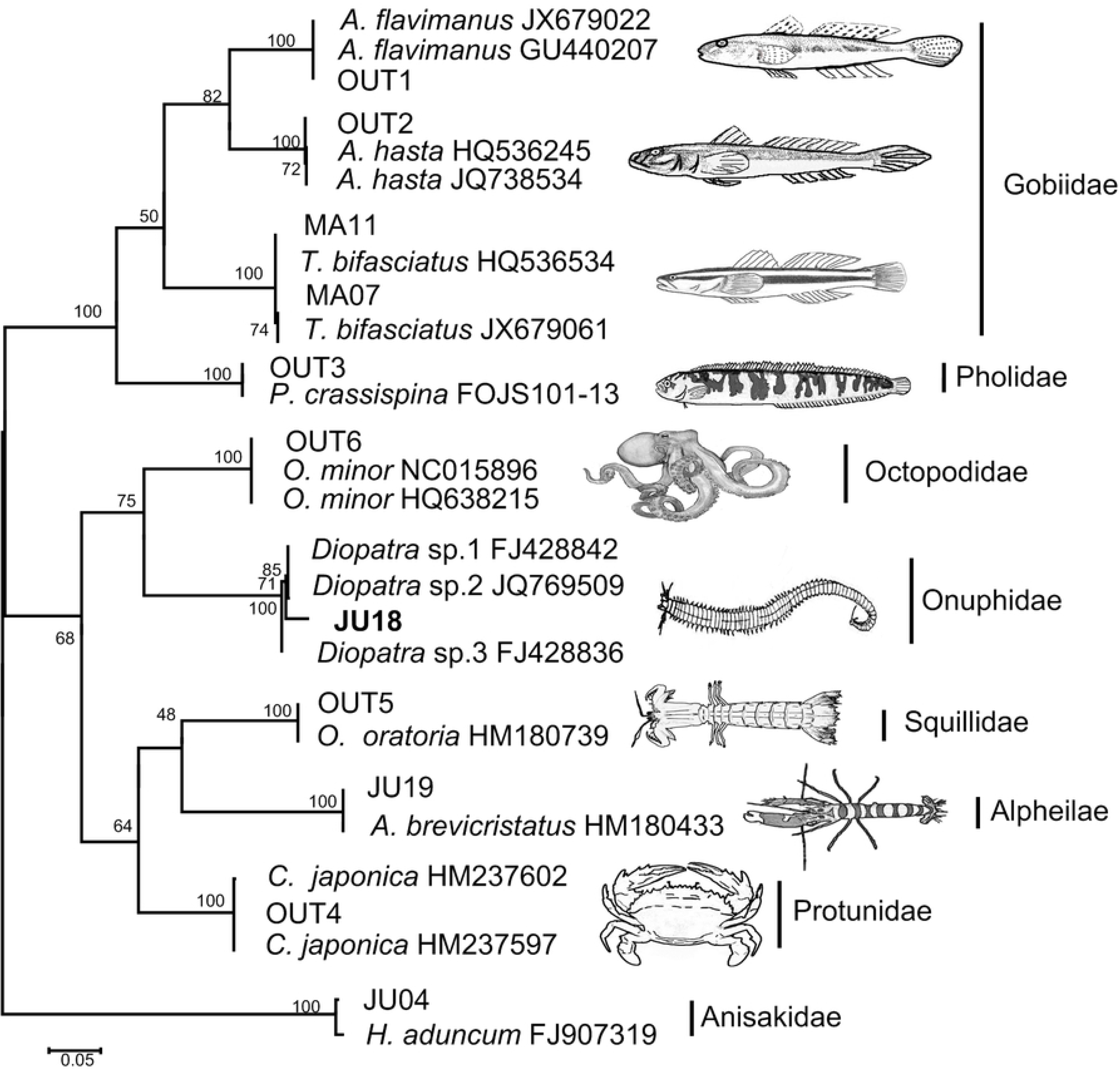
Maximum likelihood tree (ML tree) for all sequences obtained from stomach contents and the closest matches of each sequence that were downloaded from BOLD databases and GenBank, using MEGA 6 software based on K2p model

Of the 60 sequences, 30 matched with fish (50.00%, by number), 13 with crustaceans (21.66%), one with annelid (1.66%), one with nematode (1.66%) and 15 with its own species (25.00%). A total of 8 species were identified as prey of *O. minor* (Table 2). When considering the importance of prey to *O. minor* by number (%N), and ignoring the nematode and its own species as prey, the families with the highest %N were Gobiidae (61.36%), comprising *Acanthogobius flavimanus* (40.91%), *Acanthogobius hasta* (15.91%) and *Tridentiger bifasciatus* (4.55%); Protunidae (18.18%), comprising the one species *Charybdis japonica*; Pholidae (6.82%), comprising one species *Pholis crassispina*; and Squillidae (9.09%), comprising one species, *Oratosquilla oratoria.* Alpheilae and Onuphidae, accounted for same percentage, 2.27%, respectively comprising one species each, *Alpheus brevicristatus* and *Diopatra* sp. The occurrence frequency of Family of prey identified in the stomachs of *O. minor* caught in the Moon Lake was showed in Fig 3.

**Fig 3.**
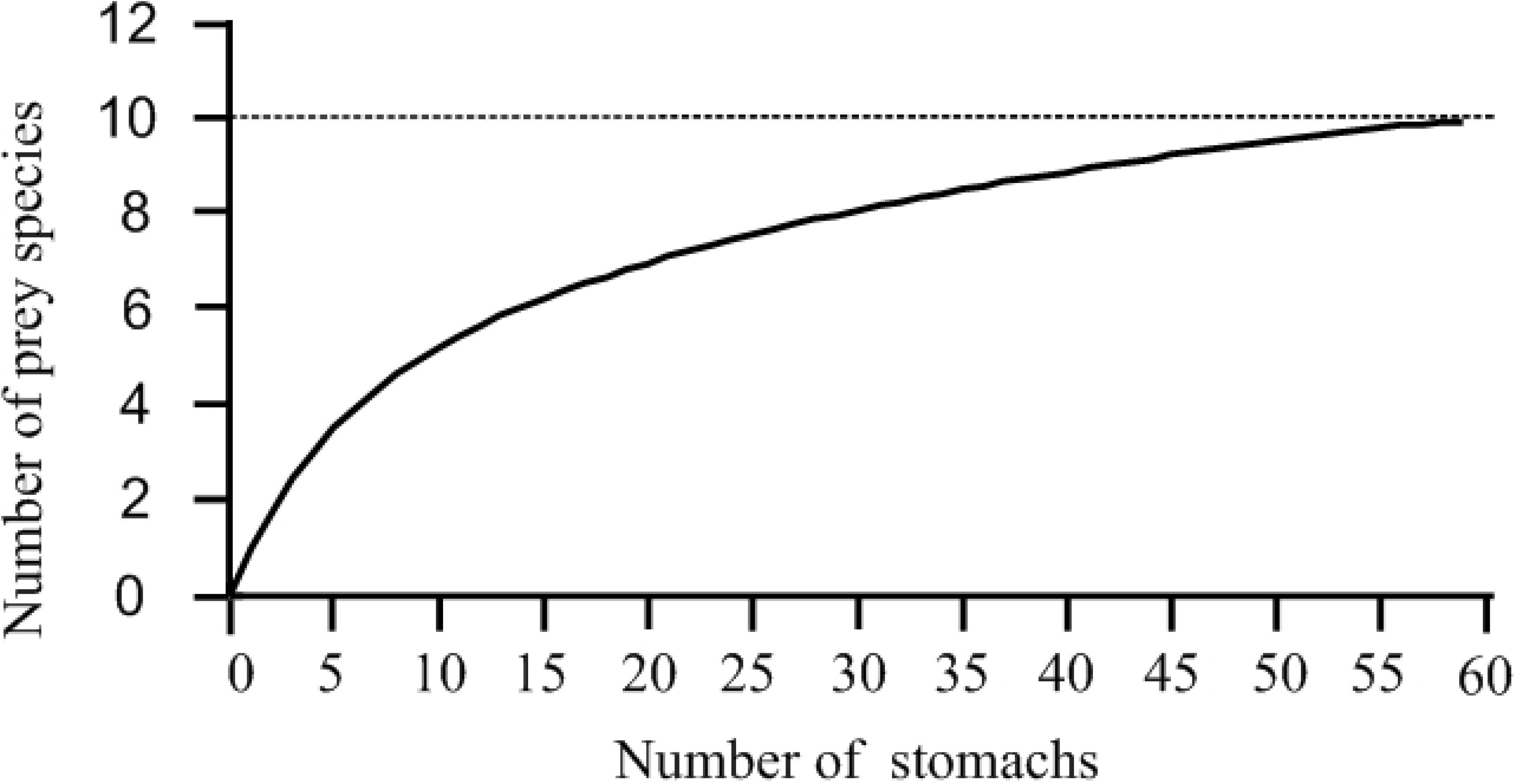
Diet of *O. minor* in Swan lake using frequency of occurrence of prey items for stomach content analysis by DNA Barcoding from April to July.

The number of prey species consumed by *O. minor* in a month was 2-5 from April to July, and in June, the prey diversity was the most abundant, with 5 species. *A. flavimanus* and *C. japonica* appeared with the highest occurring frequency during 3 months, followed by *A. hasta*. Cannibalism was found in 15 individuals, and the cannibalism occurred 2, 3, 4 and 6 cases respectively from April to July, showing an increasing trend. The cumulative prey curve approached the asymptote showing that 59 stomachs were adequate to describe the diet of this species (Fig 4).

**Fig 4.**
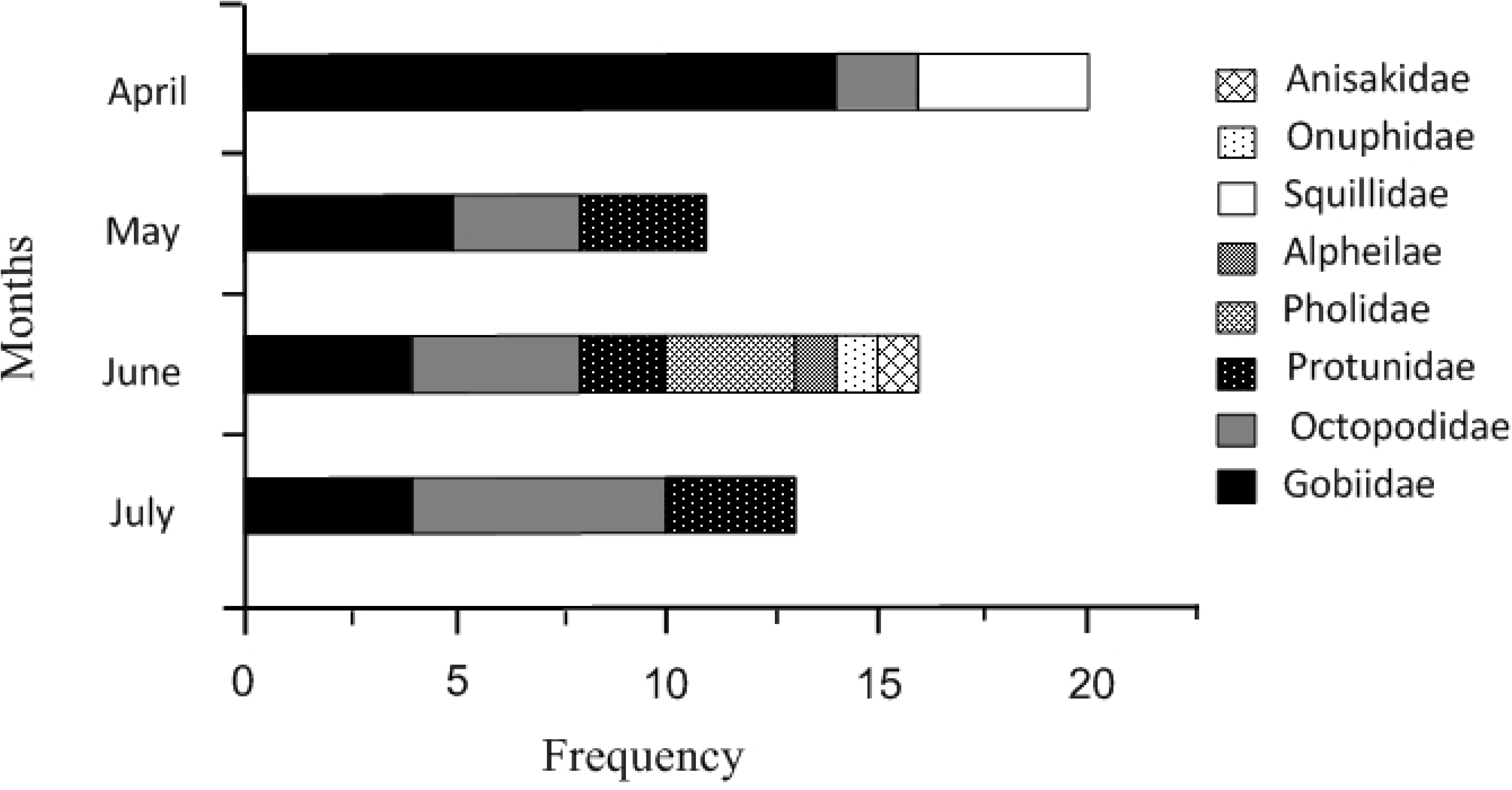
The species-accumulation cures. The slopes curve rapidly approached asymptotes, indicating that there were enough stomach content specimens collected to detect most prey species.

## Discussion

Based on our recent studies, the male octopus reaches maturity in April, meanwhile, the females are at maturing stage when the female octopus begins egg production actively, which mainly involves the synthesis of vitellus. At this stage the female octopuses use energy directly from food, with no storage reserves being transferred from organs to the gonads [34, 35]. Hence, it is unsurprising then that the feeding intensity of the female was greater than that of the male. However, the difference in feeding intensity between females and males was subsequently indistinguishable as a considerable overall feeding intensity reduction occured in both sexes caused by the listlessness of the male after mating and of the female after spawn-movement [2, 36].

One advantage of molecular methods is that when morphological methods are ineffective, sufficient DNA can be recovered by successful DNA amplifications [11, 12]. This can be achieved in poor quality samples as well because it requires only a small amount of tissue for DNA extraction. Moreover, Meusnier et al. (2008) have demonstrated that DNA barcoding could identify species with fragments as short as 100 bp with at least 90% efficiency [37]. The read lengths of the DNA fragments obtained in this study was 500-700 bp, and the relative large read lengths strengthen the accuracy of the identification. Only one clone was assigned to the genius level on the basis of their supported topographical position on the bootstrap consensus tree (Figure 2).

One intriguing discovery we found was that almost all of the stomach contents (58 out of 59) contained only one kind of prey. In consideration of stochastic sampling errors when subsampling stomach contents for DNA extraction and lack of detection of some prey species arising from low-concentration DNA to PCRs [13], duplicate subsamples with 20-40mg material each were used in this study. However, the same sequences were obtained for duplicate subsamples showed that it was unlikely that the lack of detection of diet composition was due to the procedural errors. In addition, the universal CO I primers used in this study are able to amplify CO I gene fragments from 11 invertebrate phyla and fish [30], as shown in previous studies [38, 39]. A strong dietary preference probably played a decisive role in this result. When presented with the same opportunity to different diets, *O. minor* makes a strong preference, which has been observed in other similar octopods, such as *O. maorum* [21], *O. vulgaris* [14] and *Enteroctopus dofleini* [18]. This situation has also been observed in our previous studies, where when presented with the opportunity to manila clams, the bivalve *Potamocorbula laevis* and the Asian shore crab, *O. minor* frequently preyed on the Asian shore crab and rarely on the clams. Biandolino et al. (2010) observed that only after octopus had voraciously fed on all available fish and crabs, they then fed on mussels, as the last option [24]. Ambrose (1984) found that the diet of *O. bimaculatus* in the field was influenced by food preferences, where highly preferred prey (crabs, bivalves, sedentary grazers) were rare in the field, however the most abundant species available, the snail *Tegula eiseni* Jordan, was the least preferred [20]. The results showed that although there distributed a large of mollusks, 15 molluscan species belonging to 14 families, in Moon Lake [40], octopuses did not feed on mollusks. The diet of *O. minor* in the wild was also influenced by food preferences. False negatives cannot be ruled out in this study. PCR dropout is a common phenomenon in genotyping studies because PCRs are known to amplify preferentially the DNA of a higher quality, and variable rates of DNA degradation between different prey items may have biased PCR amplification success [13]. The promise of high throughput methods in the future will only improve this approach.

The prey consisted of 8 species in this study, a relatively narrow dietary for octopus species. However, previous studies suggested that at least 12 prey species were consumed by similar octopus though examination of their middens or via morphological analyses of stomach contents [5, 6, 41]. From our analysis, the slopes of the saturation curves rapidly approached asymptotes, which indicate that sufficient stomach content specimens were collected to capture the major prey items (Fig 4). In this study we were interested in the months when the females were maturing and the dietary range wound likely be greater than reported here if the *O. minor* stomach contents had been sporadically collected from Moon Lake over a longer period [6]. Fish, including Gobiidae and Pholidae, accounted for the vast majority proportion (68.19%) of the prey. The benthic fish, Gobiidae, in the lake where the octopuses were collected, appear to be an especially important food supply for females during sexual maturation and they provided a substantial and favorite food supply for the octopods. When the preferred diet does meet the feeding needs, *O. minor* would not feed the other candidate items [17, 18, 22, 24, 32], which is a possible reason for the reduced number of types of consumed prey.

The genetic data indicate that cannibalism may be a significant feeding strategy in long-armed octopus, with long-armed octopus DNA being detected in 15 of 59 stomachs. We do not believe that the host DNA contaminated the stomach contents in these cases, as suckers and tops of arms were present in the stomachs and the host was sound with no wounds. Cannibalism in octopus has been observed in artificial culture [26, 42] and wild research [3, 43]. In this study, the number of cannibalism cases increased from April to July.

## Conclusions

The results confirm that although *O. minor* is a generalist and opportunistic predator, it has strong dietary preferences in the wild. The discovery of the feeding voracity of females during maturation provides a reference for aquaculture and artificial breeding of this commercially important species, as well as accumulating data for the rational resource exploitation of this kind animal. The DNA barcoding method was shown to be successful in unraveling the feeding habits of wild *O. minor*, which can simplify the field operation as stomach contents can be fixed by the infiltration of ethanol in the field. This method shows higher taxonomic resolution of the determination of prey items compared to traditional descriptions of stomach contents. The results provide trophic relationship information for fishery management, especially for the management of exclusive conservation reserves of *O. minor*.

## Acknowledgments

This study was supported by research grants from National Natural Science Foundation of China (Nos. 31672257; 31172058) and Fundamental Research Funds for the Central Universities(No.201822022); We are indebted to Mr. Cai B of Mashan Group Co. Ltd (Shan-Dong Province, China) and Mr. Song MP for their kind assistance in specimen collection.

